# mcSCRB-seq: sensitive and powerful single-cell RNA sequencing

**DOI:** 10.1101/188367

**Authors:** Johannes W. Bagnoli, Christoph Ziegenhain, Aleksandar Janjic, Lucas E. Wange, Beate Vieth, Swati Parekh, Johanna Geuder, Ines Hellmann, Wolfgang Enard

## Abstract

Single-cell RNA sequencing (scRNA-seq) has emerged as the central genome-wide method to characterize cellular identities and processes. While performance of scRNA-seq methods is improving, an optimum in terms of sensitivity, cost-efficiency and flexibility has not yet been reached. Among the flexible plate-based methods “Single-Cell RNA-Barcoding and Sequencing” (SCRB-seq) is one of the most sensitive and efficient ones. Based on this protocol, we systematically evaluated experimental conditions such as reverse transcriptases, reaction enhancers and PCR polymerases. We find that adding polyethylene glycol considerably increases sensitivity by enhancing cDNA synthesis. Furthermore, using Terra polymerase increases efficiency due to a more even cDNA amplification that requires less sequencing of libraries. We combined these and other improvements to a new scRNA-seq library protocol we call “molecular crowding SCRB-seq” (mcSCRB-seq), which we show to be the most sensitive and one of the most efficient and flexible scRNA-seq methods to date.

## Introduction

Whole transcriptome single-cell RNA sequencing (scRNA-seq) is a transformative tool with wide applicability to biological and biomedical questions (Wagner, Regev, and Yosef 2016). In the last few years, many new scRNA-seq protocols have been developed to overcome the challenge of isolating, reverse transcribing and amplifying the small amounts of mRNA in single cells to generate high-throughput sequencing libraries (Macaulay and Voet 2014; Kolodziejczyk et al. 2015). An idealized protocol would be able to generate one cDNA library molecule for each mRNA molecule in the cell. Such a protocol would be 100% sensitive as all mRNAs would be turned into sequenceable cDNA fragments, 100% accurate as the concentration of mRNAs would fully correlate with the number of sequenced cDNA fragments and 100% precise as the measurement error would only depend on the sampling error of sequencing reads. The lower the cost per cell for generating and sequencing a library, the more efficient the protocol would be. Furthermore, the cost per sample, i.e. the flexibility to analyze a certain number of cells from several biological samples, would also be a relevant part of the protocols efficiency. Certainly, such an optimal, one-size-fits all protocol does not and probably will never exist, as real protocols are likely to have inherent trade-offs. As a consequence, different research questions will have different optimal protocols. While many improvements have been made to scRNA-seq protocols, it is likely that further improvements are still possible. Given the importance of scRNA-seq (Regev et al. 2017), further improvements of sensitivity, efficiency and/or flexibility are also worthwhile. Each of the essential steps of scRNA-seq library preparation methods has room for improvement. First, the sensitivity of scRNA-seq methods is limited by the effectiveness of the reverse transcription and the subsequent second strand synthesis. Protocols have improved this step by optimizing enzymes, buffers and reaction volumes, resulting in conversion rates of mRNA into cDNA of 10-20% for sensitive protocols (Grün, Kester, and van Oudenaarden 2014; Svensson et al. 2017; Hashimshony et al. 2016). Second, amplification of the resulting minute amounts of cDNA leads to bias and noise when quantifying gene expression levels and hence reduce the accuracy and the precision of a scRNA-seq protocol. By incorporating random nucleotides - so called unique-molecular identifiers (UMIs) (Kivioja et al. 2012) - into the primers used for generating cDNA, amplification bias and noise can be removed by only counting cDNA fragments of a gene that have different UMIs. This increase in precision leads to a substantial increase in the power to detect differentially expressed genes in scRNA-seq protocols (Parekh et al. 2016; Ziegenhain et al. 2017). In most protocols the UMI is incorporated in the oligo-dT primer (e.g. (Soumillon et al. 2014; Jaitin et al. 2014; Macosko et al. 2015)) or in the primer used for the second-strand synthesis (e.g. (Islam et al. 2014; Arguel et al. 2017)). Therefore, the use of UMIs results in a 5’ or 3’ tag counting method and sacrifices full transcript coverage. While this can be a severe drawback when splicing and/or sequence information across the entire transcript is required, it is usually sufficient when quantifying gene expression levels to identify cell types or regulatory processes. Third, an additional and decisive advantage when reading information from the incorporated primers is that cell-specific barcodes can be incorporated during cDNA generation. This “early-barcoding” reduces costs tremendously and has allowed for the development of scRNA-seq approaches that efficiently generate libraries of tens or even hundreds of thousands of cells, especially when combined with microdroplet isolations (Macosko et al. 2015; Klein et al. 2015; Zheng et al. 2017). Hence, by incorporating early barcoding and UMIs, end-counting methods have made scRNA-seq protocols more precise and more efficient. Notably, higher amplification noise and bias still decreases the efficiency of the protocol, as more sequencing is necessary to obtain the same information.

To compare different protocols, a shared reference is needed. One such reference is a set of 92 standardized mRNAs known as ERCC spike-ins(Baker et al. 2005) that have been recently used to compare the sensitivity of different protocols mainly from published data sets (Svensson et al. 2017). While sensitivity is found to differ by more than hundred-fold, it is not clear from this comparison how the combined effects of sensitivity, measurement precision and costs per cell influence the efficiency of the protocol. Another drawback of this comparison is that ERCCs might not be fully representative for endogenous mRNAs as they are shorter, have smaller poly-A tails, do not represent the relevant concentration range with enough transcripts and are purified (Tung et al. 2017; Risso et al. 2014). Indeed, it seems that some protocols are more sensitive for ERCCs than for real mRNAs and vice versa (Ziegenhain et al. 2017). An alternative approach is to use the same cells as a shared reference and compare the cost-efficiency of protocols using power simulations (Ziegenhain et al. 2017; Vieth et al. 2017). As no standardized cells are available, this approach is currently limited to processing cells within a lab and hence to comparing only few protocols. Using this approach we found that “Single-Cell RNA-Barcoding and Sequencing” (SCRB-seq), is one of the most cost-efficient methods (Ziegenhain et al. 2017). SCRB-seq is a plate-based, early-barcoding, UMI-containing method that uses oligo-dT priming, template switching and PCR to generate amplified cDNA (Soumillon et al. 2014). Here, we set out to systematically improve the sensitivity and efficiency of SCRB-seq. Based on these evaluations, we developed molecular crowding SCRB-seq (mcSCRB-seq), a highly flexible and efficient protocol with low set-up costs that is the most sensitive scRNA-seq protocol to date.

## Design

As described above, there is the possibility and the need to improve scRNA-seq methods in terms of sensitivity and efficiency. Among plate-based methods that are efficient when processing many samples and isolating cells via FACS, SCRB-seq has been shown to be very efficient (Ziegenhain et al. 2017). Although, sensitivity and amplification bias are worse for SCRB-seq than for Smart-seq2, a methodologically similar protocol that allows for the generation of full-length scRNA-seq libraries, Smart-seq2 is less precise and more costly due to the lack of UMIs and early barcoding. As the Smart-seq2 protocol has been developed by optimizing conditions for cDNA generation (Picelli et al. 2013), this suggested that sensitivity and efficiency could also be increased for SCRB-seq. Hence, we systematically and robustly assessed how different reverse transcriptases and buffer and primer modifications impact cDNA yield from low amounts of the standardized universal human reference RNA (UHRR) (SEQC/MAQC-III Consortium 2014). We then combined the most promising improvements, in particular the addition of polyethylene glycol, and could show by sequencing the generated UHRR libraries that the new molecular crowding SCRB-seq protocol represents a 1.3-2.0 fold increase in the number of transcripts detected compared to prior versions of SCRB-seq (Soumillon et al. 2014; Ziegenhain et al. 2017). To further improve the efficiency of the new protocol by reducing the PCR amplification bias, we tested two PCR enzymes that had generated sufficient cDNA yield (KAPA HiFi and Terra) and found Terra to approximately double the library complexity at read depths below complete saturation. We then compared this optimized protocol, mcSCRB-seq, directly to a previous SCRB-seq version (Ziegenhain et al. 2017) using mouse ES cells and ERCC spike-ins. We find that it has a 50% detection probability at 2.2 ERCC transcripts, making it the most sensitive protocol among all ERCC benchmarked protocols to date. We find that it is twice as powerful in detecting differentially expressed genes than the previous SCRB-seq protocol and together with a 5-fold reduction in costs per cell and minimal hands-on time one of the most efficient and flexible scRNA-seq protocols currently available.

## Results

### A streamlined assay for cDNA yield

In order to easily quantify the effects of changes to our protocol on reverse transcription, second strand synthesis and PCR amplification, we first developed a streamlined assay to use cDNA yield as a proxy for sensitivity (Figure 1). To quantify changes to the protocol independent of biological noise, we used between 0.1 pg and 1000pg of universal human reference RNA (UHRR) as template (SEQC/MAQC-III Consortium 2014). To accommodate additions to the cDNA generation reaction more easily, we increased its volume from 2 μl (Soumillon et al. 2014) to 10 μl and confirmed that this change did not influence cDNA yield (data not shown). To quantify the cDNA yield of a single reaction, we omitted the pooling, clean-up and Exonuclease I digestion step. Instead, we heat-inactivated the reverse transcriptase and directly proceeded with PCR amplification. We measured the resulting cDNA yield by fluorometry and the cDNA length-distribution for a subset of samples with a Bioanalyzer system.

**Figure 1:**
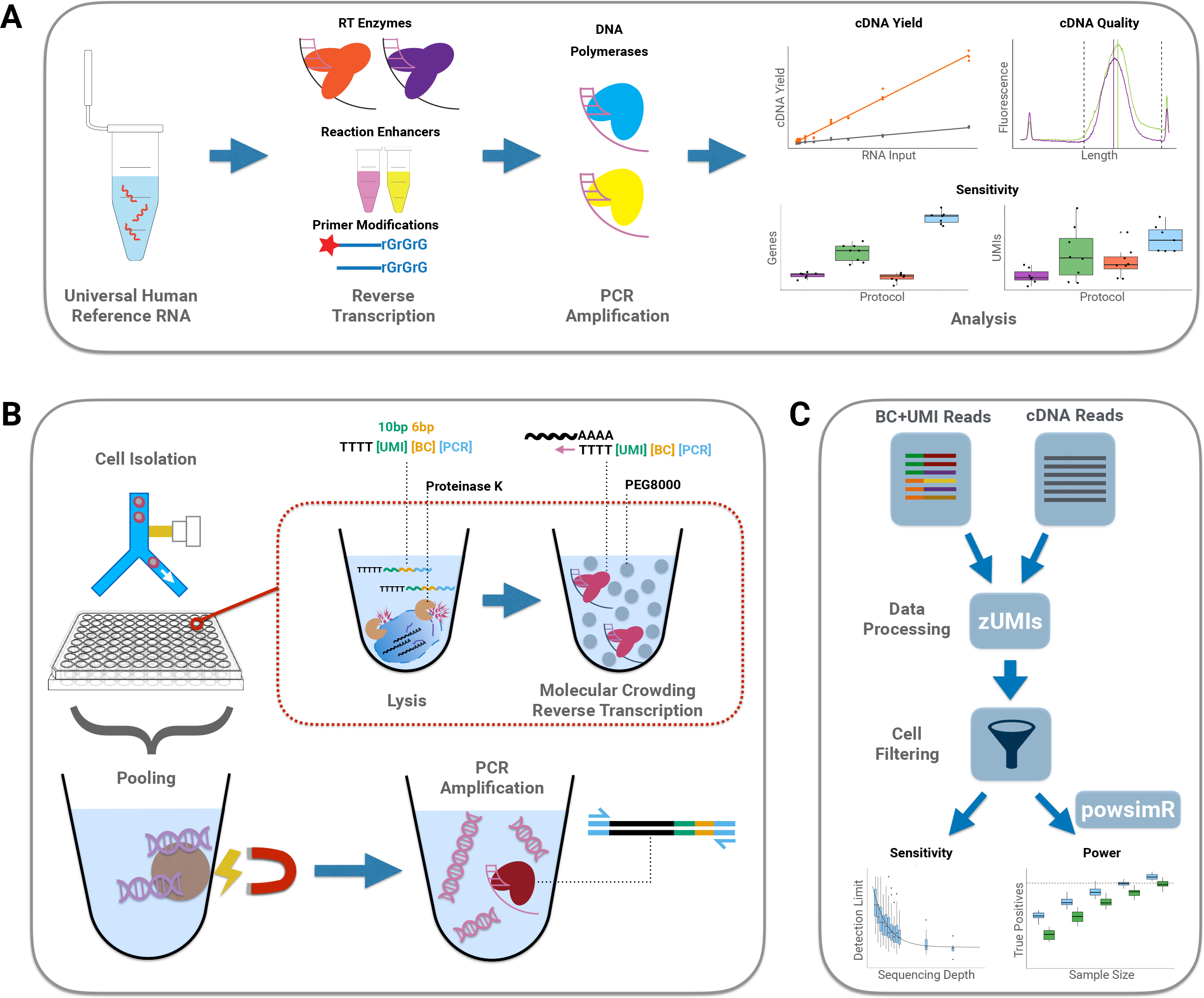
Schematic overview. **A)** Low amounts of universal human reference RNA (UHRR) were used in optimization experiments. We assessed components affecting reverse transcription and PCR amplification with respect to cDNA yield and cDNA quality and verified effects on gene and transcript sensitivity by sequencing scRNA-seq libraries to develop the mcSCRB-seq protocol. **B)** Overview of the mcSCRB-seq protocol workflow. Single cells are isolated via FACS in multiwell plates containing lysis buffer containing barcoded oligo-dT primers and Proteinase K. Reverse transcription and template-switching is carried out in the presence of PEG 8000 to induce molecular crowding. After pooling of barcoded cDNA using magnetic SPRI beads, PCR amplification using Terra polymerase is performed. **C)** Sequencing data of mcSCRB-seq libraries is processed using the zUMIs pipeline (Parekh et al. 2017). After filtering of cells, we benchmark the protocol’s performance in terms of sensitivity and power to detect differential gene expression (Vieth et al. 2017).

### cDNA yield is highest with *Maxima H*-

First, we optimized the reverse transcription reaction. In the SCRB-seq protocol, RNA is desiccated prior to reverse transcription (Soumillon et al. 2014). Our change to 10 μl reverse transcription volume allowed us to omit this step. Furthermore, we included barcoded oligo-dT primers in the lysis buffer, saving a time-consuming pipetting step in the critical phase of any scRNA-seq protocol before reverse transcription of RNA into more stable cDNA. This change resulted in a small (~10%) increase in yield (Figure 2A).

**Figure 2:**
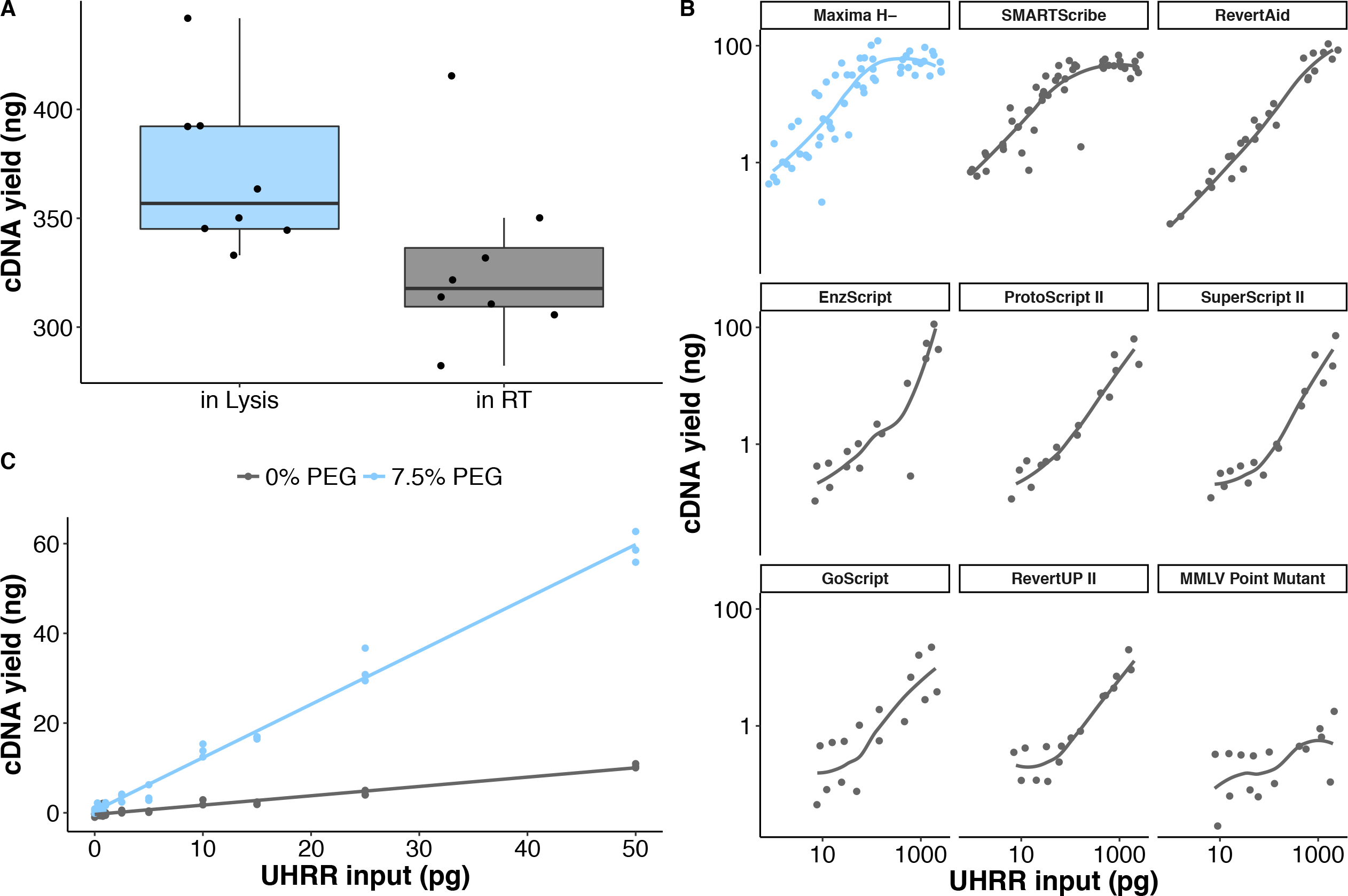
Optimizing reverse transcription sensitivity. **A)** cDNA yield (ng) after reverse transcription with oligo-dT primers already in the lysis buffer (“in Lysis”; blue) or separately added before reverse transcription (“in RT”). Each dot represents a replicate and each box represents the median and first and third quartiles. **B)** cDNA yield dependent on the absence (grey) or presence of 7.5% PEG 8000 (blue) during reverse transcription. Each dot represents a replicate. Lines represent a linear model fit of the data. **C)** cDNA yield (ng) dependent on varying UHRR input using 9 different RT enzymes. Each dot represents a replicate and fit lines were created using local regression of data points.

Similar to many scRNA-seq protocols (Ramsköld et al. 2012; Picelli et al. 2013; Islam et al. 2014; Macosko et al. 2015), our method relies on oligo-dT priming to initiate reverse transcription and a template switching reaction at the 5’ end to incorporate a priming site for preamplification. As enzyme sensitivity and processivity may be highly variable, we compared the performance of nine moloney murine leukemia virus (MMLV) reverse transcriptase enzymes that have the necessary template-switching properties. When analyzing the reaction yield in response to input amounts of RNA, Maxima H- (Thermo Fisher) and SmartScribe (Clontech) performed best (Figure 2B). Furthermore, non-MMLV reverse transcriptase enzymes (SunScript, SuperScript IV and PrimeScript II) did not yield satisfactory cDNA quality (data not shown). Notably, SuperScript II (Thermo Fisher) performed significantly worse in our experiments, contrary to other protocols (Picelli et al. 2013; Hashimshony et al. 2016).

Since pooling of cells can only occur after incorporation of cell-specific barcodes by reverse transcription, the costs for this step are a major factor in overall costs. In order to reduce enzyme costs, we showed that lowering RT enzyme to 20 units per reaction (20% reduction) does not measurably affect cDNA yield (Supplementary Figure 1A). Similarly, oligo-dT primer amounts can be reduced by 80% without repercussions (Supplementary Figure 1B). Lastly, we showed that an unblocked template-switching oligo is cheaper while retaining the same performance without primer artifacts (Supplementary Figure 1C,D).

### Molecular crowding significantly increases cDNA yield

To explore additional optimizations of the RT reaction, we evaluated additives in a previous study that had led to the increased sensitivity of the Smart-seq2 protocol (Picelli et al. 2013). Both SCRB-seq and Smart-seq2 use oligo-dT priming and template switching to generate cDNA, but surprisingly the additives that have improved cDNA yield for Smart-seq2 do not improve SCRB-seq: In our experiments, the addition of MgCl_2_ prevented the generation of full-length transcripts, while additives Betaine and Trehalose did not increase yield (Supplementary Figure 2A). What had not been explored so far for scRNA-seq protocols is adding agents such as polyethylene glycol that mimic macromolecular crowding and can drastically increase reaction rates (see (Rivas and Minton 2016) for a recent review). This effect is largely attributed to excluding solvent volume and thereby increasing the effective concentrations of reacting molecules. This can lead e.g. to more efficient ligation reactions (Zimmerman and Pheiffer 1983) and as a small reaction volume has been shown to increase the sensitivity of scRNA-seq protocols (Wu et al. 2014; Hashimshony et al. 2016; Svensson et al. 2017), we hypothesized that molecular crowding could increase the sensitivity of reverse transcription. Indeed, we observed that adding polyethylene glycol (PEG 8000) increased cDNA yield in a concentration-dependent manner (Supplementary Figure 2B). Because negative controls showed unspecific products at higher PEG-concentrations, we chose 7.5% PEG 8000 as an optimal concentration balancing yield and high specificity (Supplementary Figure 2C). With the addition of PEG 8000, yield increased dramatically, making it possible to detect RNA inputs under 1 pg (Figure 2C).

### Increases in cDNA yield translate to increased sensitivity

In order to demonstrate that our increases in cDNA yield indeed correspond to increases in sensitivity, we constructed libraries from eight replicates of 10 pg total RNA input with four protocol variants (Supplementary Table 1). Variant 1 (“Soumillon”) corresponds to the original SCRB-seq protocol (Soumillon et al. 2014), variant 2 (“Ziegenhain”) corresponds to the SCRB-seq protocol substituted with KAPA HiFi (Ziegenhain et al. 2017), variant 3 (“SmartScribe”) uses SmartScribe and KAPA HiFi, while variant 4 (“molecular crowding”) combined Maxima H-, 7.5% PEG 8000 and KAPA HiFi.

Here, the molecular crowding protocol yielded the most cDNA, while variant 1 yielded the least, confirming our systematic optimization (Figure 3A). Interestingly, variant 2 clearly outperformed variant 3, substantiating that Maxima H- is the most sensitive reverse transcriptase evaluated here. Next, we pooled all 32 libraries and sequenced 81 million reads. We used *zUMIs* (Parekh et al. 2017) to process and downsample sequencing data to one million reads per sample (Supplementary Figure 3), which has been suggested to correspond to reasonable saturation for single-cell RNA-seq experiments (Svensson et al. 2017; Ziegenhain et al. 2017). Libraries that did not obtain 1 million reads were excluded from the analysis. Taking the number of detected (>= 1 UMI) genes per sample (Figure 3B) as a first proxy for sensitivity confirmed that the molecular crowding method is the most sensitive protocol (p = 7 × 10^−7^, Welch Two Sample t-test, compared to variant 2) with 7,898 genes on average, while variants 1-3 detected only 3,938, 5,542, 3,805 genes, respectively. As our data contained UMIs, we could then use the number of total detected molecules per sample as a second measure of sensitivity (Figure 3C). Although more variable, this corroborated our findings on detected genes. Next, we asked whether the increase in sensitivity translates not only in more detected genes but also in more reproducible detection of genes. For this, we calculated the dropout probabilities of genes, excluding stochastically detected genes (<0.2 UMIs mean expression) (Lun, Bach, and Marioni 2016). Confirming our previous findings, molecular crowding markedly improved detection rates. Clearly visible, genes had lower overall dropout probabilities and a significantly larger number of genes was detected in all samples (Figure 3D).

**Figure 3:**
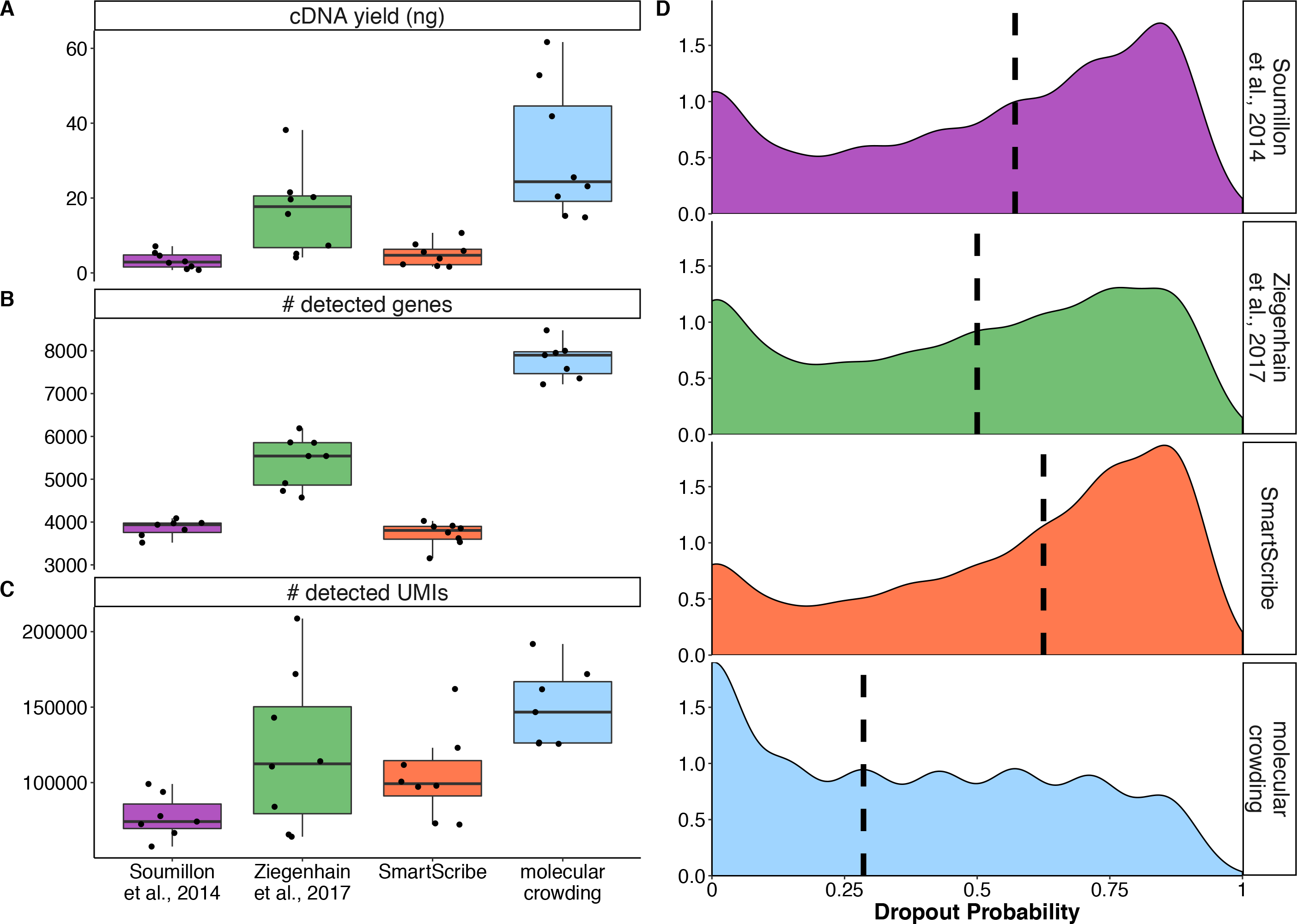
Molecular crowding increases sensitivity. **A-D)** RNA-seq libraries were generated from 10 pg of UHRR using four protocol variants (see Supplementary Table 1). **A)** cDNA yield (ng) after PCR amplification per method. Each dot represents a replicate and each box represents the median and first and third quartiles per method. **B)** Number of genes detected (>1 UMI) per replicate at a sequencing depth of one million raw reads. Each dot represents a replicate and each box represents the median and first and third quartiles per method. **C)** Number of unique molecular identifiers per replicate at a sequencing depth of one million raw reads. Each dot represents a replicate and each box represents the median and first and third quartiles per method. **D)** Gene dropout probability (0-1) over all replicates for each method at a sequencing depth of one million raw reads.

### Terra polymerase retains library complexity during PCR

Single-cell RNA sequencing methods rely on amplifying very low amounts of input material. It is well established that noise and bias may be introduced during library PCR, depending on the number of cycles, reaction conditions and polymerases (Parekh et al. 2016; Quail et al. 2012). While UMIs can largely correct the effects of noise and bias, it still requires more reads to reach the same information, resulting in a higher efficiency of scRNA-seq methods that have less amplification bias (Ziegenhain et al. 2017; Sasagawa et al. 2017). To optimize PCR conditions, we first evaluated twelve polymerases for cDNA yield. Three polymerases (KAPA HiFi, SeqAmp and Terra) yielded significantly more amplified cDNA after 18 PCR cycles (Supplementary Figure 4A) than the Advantage2 enzyme that was used in the original protocol (Soumillon et al. 2014). We disregarded SeqAmp because of a decreased median length of the amplified cDNA molecules (Supplementary Figure 4B) and compared amplification bias and noise of the KAPA and Terra polymerases by generating libraries from single mouse embryonic stem cells (mESCs) using our optimized molecular crowding protocol to generate cDNA. We pooled cDNA from 32 cells and amplified cDNA using either KAPA or Terra polymerase. After sequencing both library pools, we processed the data and downsampled each transcriptome to one million raw reads to exclude bias from varying coverage (Parekh et al. 2017). Taking the number of detected UMIs per cell as a measure, we found that PCR amplification using Terra yielded twice as much library complexity than using KAPA (Supplementary Figure 4C). Thus, we chose the Terra polymerase for the mcSCRB-seq protocol in order to retain as much as possible of the initial transcriptome complexity through preamplification. Importantly, the higher yield of the molecular crowding reverse transcription allowed us to reduce the number of PCR cycles, thereby further reducing amplification bias (Parekh et al. 2016).

### mcSCRB-seq increases sensitivity 2.5-fold over previous SCRB-seq protocols

In order to assess the improvements of the entire molecular crowding SCRB-seq (mcSCRB-seq) protocol in comparison to the previously benchmarked SCRB-seq protocol used in Ziegenhain et al. (Supplementary Table 2), we generated libraries from mouse ES cells (mESCs) including spiked-in ERCCs (Baker et al. 2005). We used a single sample of mESCs and sorted two plates containing 96 and 48 cells for each method. Libraries were prepared on the same day and multiplexed for sequencing in order to avoid batch effects. Following sequencing, we filtered cells by excluding doublets identified from the distribution of per-cell total UMI counts (Ziegenhain et al. 2017). Furthermore we discarded broken cells and failed libraries by inspecting nearest-neighbor correlation of gene expression values (Petropoulos et al. 2016), yielding 249 high-quality libraries (Supplementary Figure 5). The percentage (median 88%) and distribution of mapped reads (~50% in exons) was very similar in all four libraries (Supplementary Figure 6). To assess sensitivity and library complexity relative to sequencing depth we downsampled reads to fixed depths using the *zUMIs* pipeline (Parekh et al. 2017). Already at low depths mcSCRB-seq clearly outperformed SCRB-seq and detected 2.5 times as many UMIs per cell at sequencing depths above 200,000 reads (Figure 4A). At 500,000 reads mcSCRB-seq detected 50,969 UMIs that corresponded to 5,866 different genes, 1,000 more than SCRB-seq (Supplementary Figure 7). Interestingly, libraries were not sequenced to saturation at one million reads as the number of detected UMIs still increased at this depth (Supplementary Figure 7B). In order to judge the absolute sensitivity of mcSCRB-seq, we used ERCC spike-ins to estimate the RNA content per cell by dividing the number of detected transcriptomic UMIs by the fraction of ERCC UMIs detected from the total number of spiked-in ERRC molecules (Supplementary Figure 8). Fitting with previous reports (Islam et al. 2014), the median mRNA content of our mouse ES cells was 227,467 molecules. Using this estimate, we could then convert the number of transcriptomic UMIs detected to the fraction of the cellular mRNAs that was observed at different sequencing depths (Figure 4B). At a depth of 2 million reads per cell mcSCRB-seq could detect above 50% of the cellular mRNA content, 2.5-fold exceeding the 20% sensitivity of SCRB-seq and the estimated sensitivity of previous protocols (Grün, Kester, and van Oudenaarden 2014). As expected, this higher sensitivity of mcSCRB-seq also lead to a larger number of genes detected in all cells (Supplementary Figure 9A) and to a more reliable detection of genes, i.e. a lower dropout rate, within cells (Supplementary Figure 9B). Congruent with the previous comparison of Terra and KAPA polymerase for amplifying cDNA, mcSCRB-seq showed a more uniform amplification, as the extra-poisson variability of reads per genes was lower in the mcSCRB-seq protocol (Supplementary Figure 9C). Although both methods use UMIs to remove PCR bias (Supplementary Figure 9D), the reduction of the preamplification variance leads to higher information content at the same sequencing depth.

**Figure 4:**
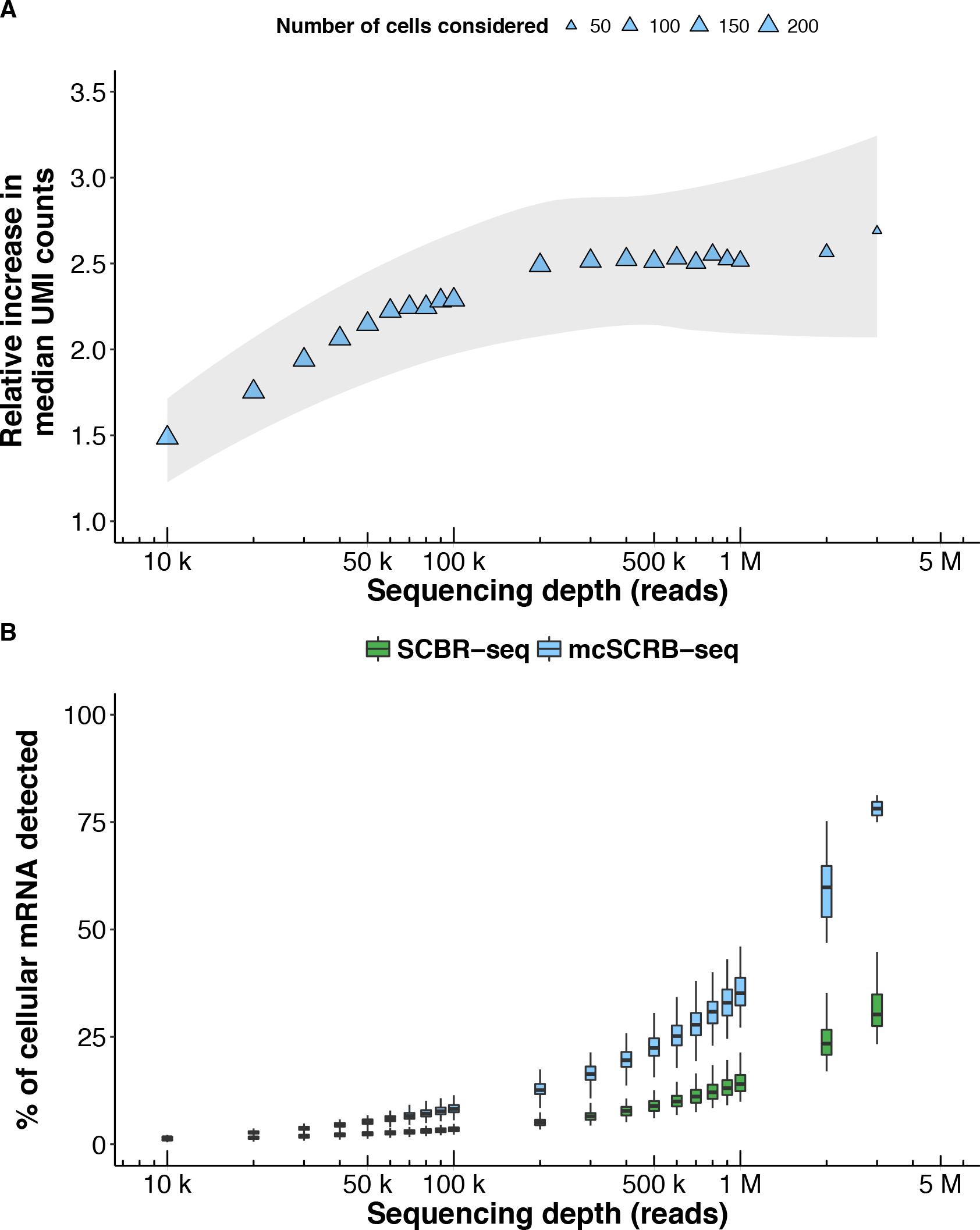
mcSCRB-seq detects large fractions of the cellular transcriptome of mESCs. **A)** Relative increase in the median of detected UMIs dependent on raw sequencing depth (reads) using mcSCRB-seq compared to SCRB-seq. Each symbol represents the median over all cells at the given sequencing depth. The size of symbols depicts the number of cells that were considered to calculate the median. The 95% confidence interval of a local regression model is depicted by the shaded area. **B)** Percentage of cellular mRNA content (see Supplementary Figure 8) that can be detected with SCRB-seq (green) or mcSCRB-seq (blue) dependent on the sequencing depth (reads). Each box represents the median and first and third quartiles per sequencing depth and method.

### mcSCRB-seq is the most sensitive protocol as benchmarked by ERCCs

The widespread use of ERCC spike-ins allows us to estimate and compare the absolute sensitivity across many scRNA-seq protocols using published data (Svensson et al. 2017). As in Svensson et al., we used a binomial logistic regression to estimate the number of ERCC transcripts that are needed on average to reach a 50% detection probability. mcSCRB-seq reached this threshold with 2.2 molecules when ERCCs are sequenced to saturation (Figure 5B). Comparing this to a total of 26 estimates for 20 protocols obtained from the two major protocol comparisons (Svensson et al. 2017; Ziegenhain et al. 2017) as well as additional relevant protocols, such as Quartz-seq2 (Sasagawa et al. 2017) and the 10x Genomics Chromium chemistry (Zheng et al. 2017), mcSCRB-seq indeed shows the highest sensitivity among all protocols compared to date (Figure 5C).

**Figure 5:**
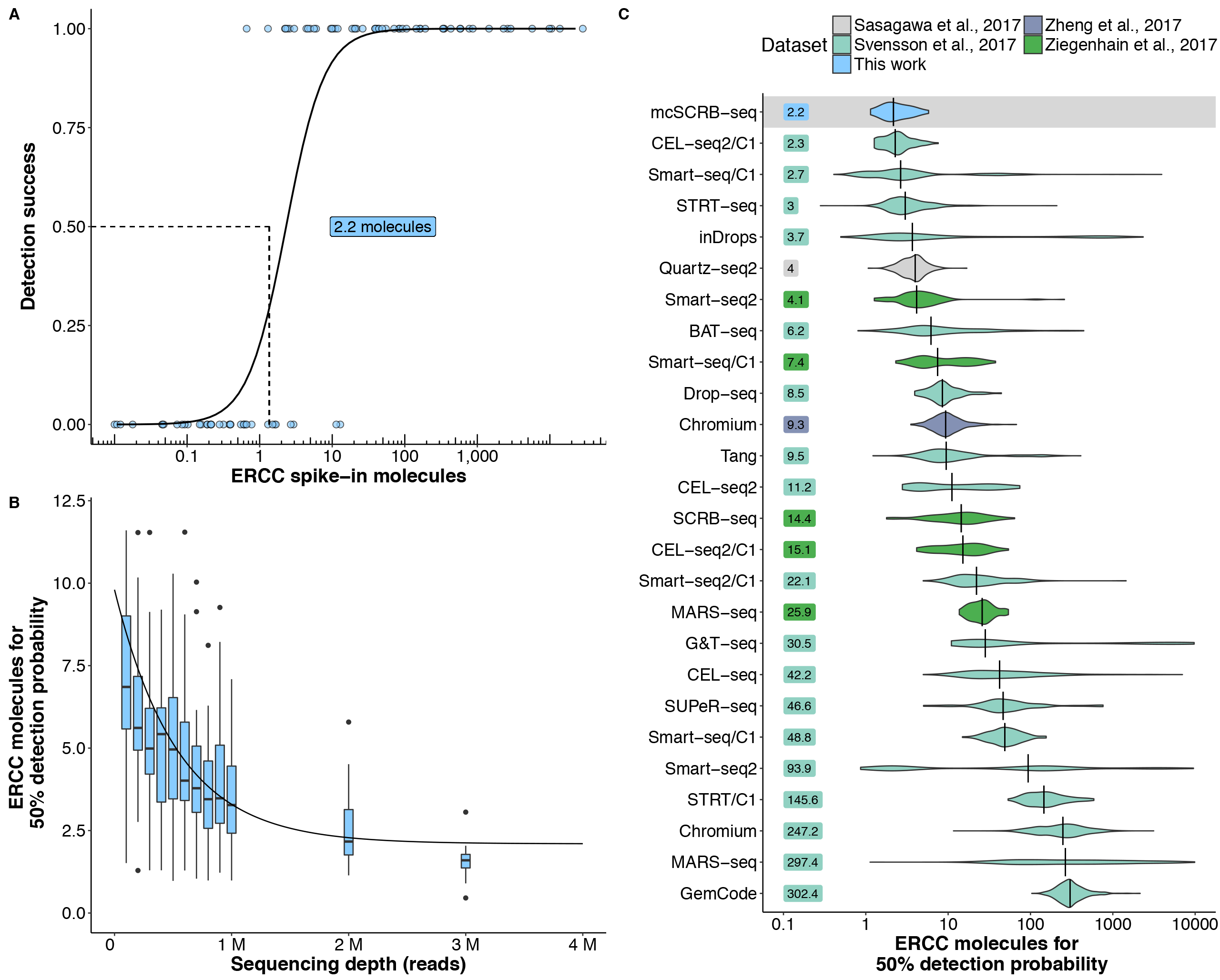
mcSCRB-seq is the most sensitive protocol using ERCC spike-ins. **A-C)** Detection of ERCC spike-in transcripts was modeled using a binomial logistic regression relative to the input molecule number. **A)** Shown is the detection of the 92 ERCC transcripts in an average cell processed with mcSCRB-seq at 2 million reads coverage. Points and solid line represent the ERCC genes with their logistic regression model. Dashed lines and label indicate the number of ERCC molecules required for a detection probability of 50%. **B)** Number of ERCC molecules required for 50% detection probability dependent on the sequencing depth (reads) for mcSCRB-seq. Each each box represents the median, first and third quartiles of cells per sequencing depth with dots marking outliers. A non-linear asymptotic fit is depicted as a solid black line. **C)** Number of ERCC molecules required for 50% detection probability for various library preparation protocols. Per-cell distributions are shown using violin plots, vertical lines and labels depict the median per protocol.

### mcSCRB-seq combines high power, fast processing and low costs

The more sensitive a scRNA-seq method is, the greater its power, as long as it utilizes UMIs that allow it to remove amplification noise (Ziegenhain et al. 2017). As expected, when using simulations (Vieth et al. 2017) to compare the power of mcSCRB-seq and SCRB-seq to detect differentially expressed genes at a sequencing depth of 500,000 reads per cell, we find that mcSCRB-seq requires approximately half as many cells to reach the same power as SCRB-seq (Figure 6A and B). Furthermore, expression levels differed much less between batches of mcSCRB-seq libraries (Supplementary Figure 10A), indicating that it might be a more robust protocol than SCRB-seq. Though, many more batches across labs and conditions would be required to systematically compare robustness of protocols. We did not find biased expression levels related to GC content or transcript lengths in SCRB-seq or mcSCRB-seq (Supplementary Figure 10B,C), unlike what has been found in other protocols (Phipson, Zappia, and Oshlack 2017). In our recent comparison, SCRB-seq was already among the most cost-efficient scRNA-seq protocols, i.e. the minimal costs for generating and sequencing scRNA-seq libraries from enough cells to reach 80% power were similarly low for SCRB-seq, MARS-seq and Drop-seq (Ziegenhain et al. 2017). mcSCRB-seq halves the cost by doubling the power (Figure 6A). Furthermore, our additional optimizations reduced costs from 2 € per cell for SCRB-seq to less than 0.6 € per cell in a 96-well format and to less than 0.4 € per cell in a 384-well format (Figure 6B, Supplementary Table 3). Moreover, owing to an optimized workflow, we could reduce the library preparation time to one working day with minimal hands-on time (Figure 6C, Supplementary Table 4).

**Figure 6:**
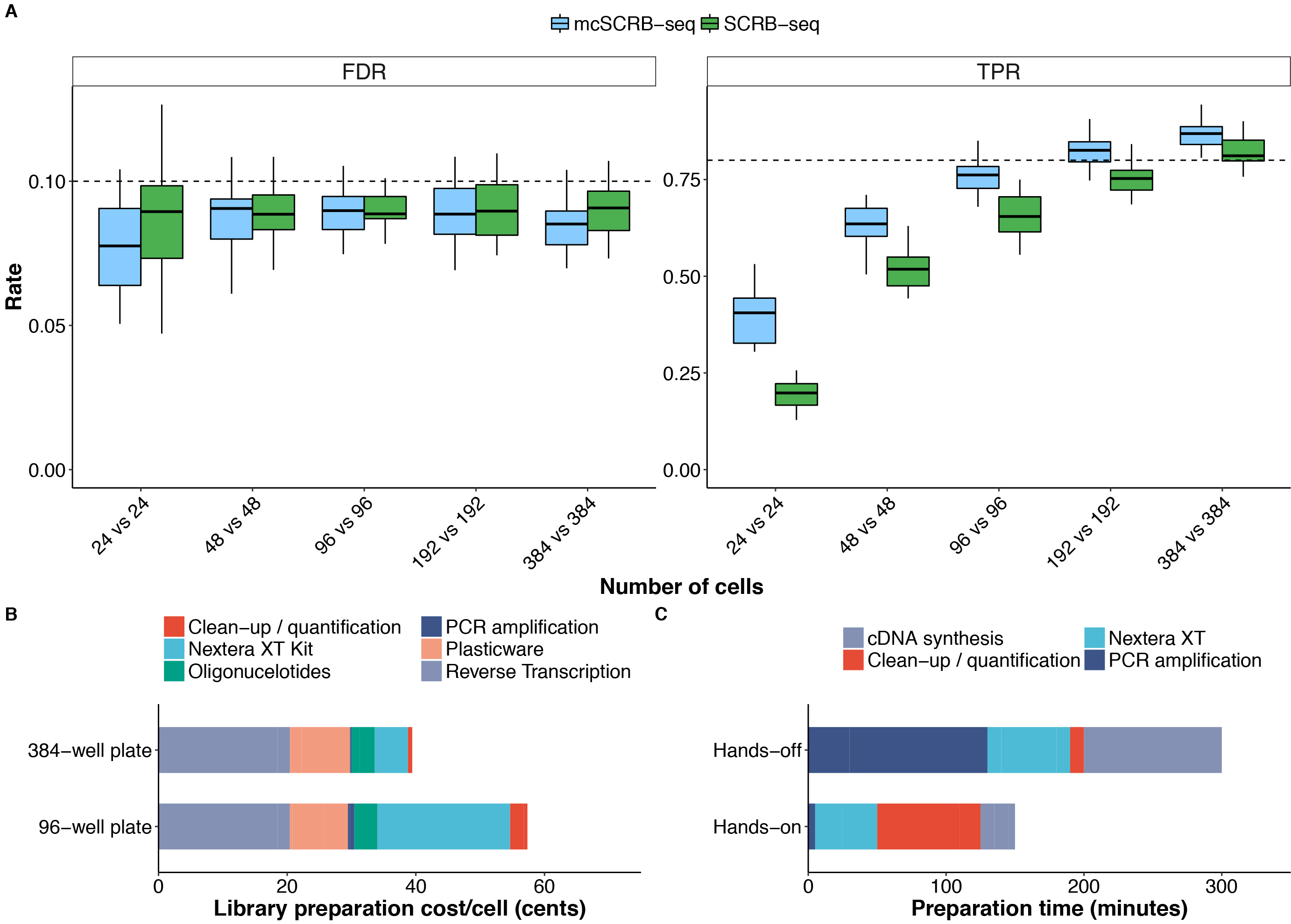
mcSCRB-seq is highly powerful and efficient. **A)** Power simulations were performed using the powsimR package (Vieth et al. 2017) from empirical parameters estimated at 500,000 raw reads per cell. For SCRB-seq and mcSCRB-seq, we simulated *n*-cell two-group differential gene expression experiments with 10% differentially expressed genes. Shown are true positive rate (“TPR”) and false discovery rate (“FDR”) for sample sizes n = 24, n = 48, n = 96, n = 192 and n = 384 per group. Boxplots represent the median and first and third quartiles of 25 simulations. Dashed lines indicate the desired nominal levels of TPR and FDR. **B)** Library preparation costs per cell were calculated for 96-well or 384-well scenarios. Colors indicate the consumable type. (see Supplementary Table 3) **C)** Library preparation time for one 96-well plate of mcSCRB-seq libraries was measured for bench times (“Hands-on”) and incubation times (“Hands-off”). Colors indicate the library preparation step. The total time was 7.5 hours. (see Supplementary Table 4)

Taken together, we show that mcSCRB-seq considerably increases sensitivity due to the addition of PEG and the reduced amplification bias of the Terra polymerase, requiring half as many cells than SCRB-seq to reach the same power. In addition, optimized reagents and workflows reduce costs per cell up to five-fold, leading to an up to ten-fold increase in cost-efficiency compared to SCRB-seq. Hence, mcSCRB-seq is not only the most sensitive protocol when benchmarked using ERCCs, it is probably also one of the most cost-efficient protocols currently available.

## Discussion

Here, we have presented a systematic optimization of SCRB-seq, a plate-based, 3’ tagging scRNA-seq protocol, resulting in mcSCRB-seq, one of the most sensitive and efficient protocols currently available. This has implications for understanding, further developing and applying scRNA-seq protocols.

Maybe the most important insight of our systematic optimization of cDNA yield is that adding polyethylene glycol (PEG) can strongly enhance cDNA generation from small amounts of starting material. A likely mechanisms is that the macromolecule PEG reduces the accessible volume of the reaction, hence effectively increasing the concentrations and in turn reaction rates (Ellis 2001). This effect of macromolecular crowding has been used to increase the rate of DNA ligations (Zimmerman and Pheiffer 1983) and is also thought to occur in cells where 20-30% of the volume is occupied by macromolecules (Han and Herzfeld 1993). If PEG indeed speeds up the reaction, it could increase cDNA yield by shifting the balance from enzymatic and chemical mRNA degradation towards reverse transcription. Such a scenario would be compatible with the findings that speeding up the reaction by providing oligo-dT primers already in the lysis buffer also increases cDNA yield (Figure 2A). In any case, it will be interesting to see whether the addition of PEG might increase the sensitivity of other scRNA-seq protocols. However, the interaction of reagents seems complex, as improvements across protocols are not necessarily transferable. In our case, altering MgCl_2_, Betaine and Trehalose concentrations that led to improvements for the sensitive Smart-seq2 protocol (Picelli et al. 2013) did not enhance cDNA yield, although both protocols are very similar as they generate cDNA by oligo-dT priming, template switching and PCR amplification. Given this complex interaction of primers, enzymes and reaction conditions, it is likely that sensitivity can be further optimized for many protocols, although it could require many experiments to find the optimum.

While the efficiency of cDNA generation is probably the rate limiting step for sensitivity, it is important to realize that biased cDNA amplification also can strongly impact sensitivity when libraries are not sequenced to saturation. That this bias is substantial can be visualized by plotting the coefficient of variance (CV) against the mean expression level for each gene or similarly by calculating the variability that is added to on top of the expected poisson sampling variability (Supplementary Figure 9C,D). When considering read counts, there is much more extra-poisson variability than when considering UMIs. This shows that most of the extra-poisson variability is due to technical amplification bias and not due to biological variation as generally known for scRNA-seq protocols (Ziegenhain et al. 2017). Importantly, the range of this bias among genes is substantial, ranging from genes that are hardly amplified to others that are amplified very strongly. While this bias can be corrected for by using UMIs, it still uses up a substantial amount of sequencing reads, limiting the sensitivity when libraries are not sequenced to saturation and making the method less cost-efficient across all sequencing depths. Here, we have optimized this property by using Terra polymerase for cDNA amplification that produces slightly less yield, but substantially less bias than KAPA polymerase. This property of the Terra polymerase has also been used to make the Quartz-seq2 protocol efficient (Sasagawa et al. 2017). How the biased amplification in combination with the sensitivity of cDNA generation can be optimized to generate more sensitive and efficient scRNA-seq methods has not been explored much for most methods and has considerable potential for optimizing scRNA-seq protocols.

Which implications does mcSCRB-seq have for applying scRNA-seq, i.e. under what circumstances should one consider using mcSCRB-seq over the >40 other scRNA-seq protocols? We have shown by using standardized RNA and parallelly isolated and processed mESCs that mcSCRB-seq clearly outperforms a previous version of SCRB-seq and even more so the original version of SCRB-seq in terms of sensitivity and costs. The increased sensitivity leads to almost a 2-fold increase in power, i.e. half as many cells need to be analyzed to have the same power to detect differentially expressed genes. Optimized reagents reduce the costs per cell from ~2 € to 0.4 € and an optimized procedure reduces the preparation time of libraries to a working day. As SCRB-seq, together with MARS-seq and Drop-seq, was already among the most efficient protocols in our previous power comparison, mcSCRB-seq is likely to be up to ten-times more cost-efficient than SCRB-seq (Ziegenhain et al. 2017). Note that the absolute sensitivity of detected genes or transcripts for mESCs is higher for cells processed by SCRB-seq in Ziegenhain et al. than the mESCs processed by exactly the same SCRB-seq protocol here. This suggests that subtle differences in cell line, passage number, culture conditions and/or cell isolation can lead to considerable differences in mRNA content per cell, cautioning comparisons done even with the same cell types across time and laboratories. In other words, using freshly cultured cells as a benchmark to compare scRNA-seq protocols is more limited than one might expect. A possible alternative could be standardized frozen cells distributed from a central resource, but such a standard is currently not available. An alternative is standardized purified RNA from a complex source, like the UHRR RNA used here. Obviously, such a standard has limitations as it does not allow compare methods for factors acting before cell lysis and furthermore it has not been widely used to benchmark scRNA-seq protocols so far. A standard that has been extensively used and has the additional advantage of known mRNA concentrations are the ERCC spike-ins. While it is unclear to what extent ERCCs are fully representative of normal mRNAs, they are currently the only possibility to compare a wide range of protocols within and across laboratories (Svensson et al. 2017). Calculating the 50% detection probability of ERCCs, as done by Svensson et al., we find that mcSCRB-seq is indeed the protocol with the highest median sensitivity for ERCCs. This is in line with our finding that mcSCRB-seq is 2.5 fold more sensitive than SCRB-seq, which was already very sensitive when compared to other methods (Ziegenhain et al. 2017). Importantly, as we have argued before (Ziegenhain et al. 2017), efficiency is what matters in the end for most experiments, i.e. the costs to generate and sequence scRNA-seq libraries at given amount of power. Unfortunately, 92 ERCC genes are not sufficient for reasonable power simulations, so it is currently not possible to quantitatively compare the cost-efficiency across many protocols. However, it is clear that the protocols that are almost as sensitive as mcSCRB-seq and use Fluidigm’s C1 are more than ten-times more expensive per cell than mcSCRB-seq (Ziegenhain et al. 2017) and hence clearly less efficient. Among other prominent plate-based methods, like CEL-seq2, Quartz-seq2 or STRT-seq, mcSCRB-seq is the most sensitive one and with 0.4 € per cell possibly even the cheapest plate-based method. Droplet-based methods like inDrop (Klein et al. 2015), Drop-seq (Macosko et al. 2015) or the 10x Genomics system (Zheng et al. 2017) and other random distributing methods like sci-RNA-seq (Cao et al. 2017), SPLiT-seq (Rosenberg et al. 2017) or Seq-well (Gierahn et al. 2017) are cheaper per cell, at least when many cells per sample are processed. It is currently unclear in which context which scRNA-seq protocols will turn out to be optimal in terms of cost-efficiency, cell isolation procedures and additional features like cell imaging, multi-omics measurements and/or compatibility with fixatives. Better standards like centrally distributed cells or at least spike-ins that better represent endogenous mRNAs would certainly be very helpful for more quantitative comparisons. In any case, it seems very likely that sensitive and flexible plate-based methods in combination with FACS-based cell isolation will remain an important part of the scRNA-seq portfolio, certainly as long as sequencing costs do not drop by an order of magnitude. Among those plate-based methods, mcSCRB-seq is one of the most sensitive and most efficient. It is fast, flexible, easy to set-up and hence could be a valuable methodological addition for many laboratories.

### Limitations

As mcSCRB-seq is a plate-based method, one major limitation is that a FACS is required to efficiently isolate cells and the plate-based format can not easily reach the throughput of thousands of cells per sample as possible for droplet-based or combinatorial indexing methods (Macosko et al. 2015; Klein et al. 2015; Zheng et al. 2017; Cao et al. 2017; Rosenberg et al. 2017). Furthermore, it is a 3’ counting method and hence is not well suited to analyse splicing patterns and sequence variants present further upstream, as possible with full-length methods like Smart-seq2 (Picelli et al. 2013). mcSCRB-seq has not been tested for isolating nuclei or for multi-omic measurements, but this current limitation might well be overcome in the future.

## Methods

### Optimization experiments

For all optimization experiments, universal human reference RNA (UHRR; Agilent) was utilized to exclude biological variability. Unless otherwise noted, 1 ng of UHRR was used as input per replicate. Additionally, Proteinase K digestion and desiccation were not necessary prior to reverse transcription. In order to accommodate all reagents into the reaction, the total volume for reverse transcription was increased to 10 μl. While all concentrations were kept the same, we added the same total amount of reverse transcriptase (25 U), with its concentration thus lowering from 12.5 U/μl to 2.5 U/μl. After reverse transcription, no pooling was performed, rather preamplification was done per replicate. For each sample, we measured the cDNA concentration using the Quant-iT PicoGreen dsDNA Assay Kit (Thermo Fisher).

### Comparison of reverse transcriptases

Nine reverse transcriptases, Maxima H- (Thermo Fisher), SMARTScribe (Clontech), Revert Aid (Thermo Fisher), EnzScript (Biozym), ProtoScript II (New England Biolabs), Superscript II (Thermo Fisher), GoScript (Promega), Revert UP II (Biozym), M-MLV Point Mutant (Promega), were compared to determine which enzyme resulted in the largest cDNA yield. Several dilutions ranging from 10 to 1000 pg of universal human reference RNA (UHRR; Agilent) were used as input into the RT reactions.

RT reactions contained final concentrations of 1× M-MuLV reaction buffer (NEB), 1 mM dNTPs (Thermo Fisher), 1 μM E3V6NEXT barcoded oligo-dT primer (IDT), and 1 μM E5V6NEXT template-switching oligo (IDT). For reverse transcriptases with unknown buffer conditions, the provided proprietary buffers were used. Reverse transcriptases were added for a final amount of 25 U per reaction.

### Effect of reaction enhancers

In order to improve the efficiency of the RT, we tested the addition of reaction enhancers, including MgCl_2_, betaine, trehalose, and polyethylene glycol (PEG 8000). The final reaction volume of 10 μL was maintained by adjusting the volume of H_2_O.

For this, we added increasing concentrations of MgCl_2_ (3, 6, 9, and 12 mM; Sigma-Aldrich) in the RT buffer in presence or absence of 1 M betaine (Sigma-Aldrich). Furthermore, the addition of 1 M betaine and 0.6 M trehalose (Sigma-Aldrich) was compared to the standard RT protocol. Lastly, increasing concentrations of PEG 8000 (0, 3, 6, 9, 12, 15 % W/V) were also used.

### Comparison of PCR DNA polymerases

The following twelve DNA polymerases were evaluated in preamplification: KAPA HiFi HotStart (KAPA Biosystems), SeqAmp (Clontech), Terra direct (Clontech), Platinum SuperFi (Thermo Fisher), Precisor (Biocat), Advantage2 (Clontech), AccuPrime Taq (Invitrogen), Phusion Flash (Thermo Fisher), AccuStart (QuantaBio), PicoMaxx (Agilent), FideliTaq (Affymetrix), Q5 (New England Biolabs). For each enzyme, at least three replicates of 1 ng UHRR were reverse transcribed using the optimized molecular crowding reverse transcription in 10 μl reactions. Optimal concentrations for dNTPs, reaction buffer, stabilizers, and enzyme were determined using manufacturer’s recommendations. For all amplification reactions, we used the original SCRB-seq PCR cycling conditions (Soumillon et al. 2014).

### Cell culture of mouse embryonic stem cells

J1 (Li, Bestor, and Jaenisch 1992) and JM8 (Pettitt et al. 2009) mouse embryonic stem cells were cultured under feeder-free conditions on gelatine-coated dishes in high-glucose Dulbecco’s modified Eagle’s medium (Thermo Fisher) supplemented with 15% fetal bovine serum (FBS, Thermo Fisher), 100 U/ml penicillin, 100 μ g/ml streptomycin (Thermo Fisher), 2 mM L-glutamine (Thermo Fisher), 1× MEM non-essential amino acids (NEAA, Thermo Fisher), 0.1 mM β-mercaptoethanol (Thermo Fisher), 1000 U/ml recombinant mouse LIF (Merck Millipore) and 2i (1 μM PD032591 and 3 μM CHIR99021 (Sigma-Aldrich)). mESCs were routinely passaged using 0.25% trypsin (Thermo Fisher).

mESC cultures were confirmed to be free of mycoplasma contamination by a PCR-based test (Young et al. 2010).

### SCRB-seq cDNA synthesis

Cells were dissociated using trypsin and resuspended in 100 μL of RNAprotect Cell Reagent (Qiagen) per 100 000 cells. Directly prior to FACS sorting, the cell suspension was diluted with PBS (Gibco). Single cells were sorted into 96-well DNA LoBind plates (Eppendorf) containing lysis buffer using a Sony SH800 sorter (Sony Biotechnology; 100 μm chip) in “Single Cell (3 Drops)” purity. Lysis buffer consisted of a 1:500 dilution of Phusion HF buffer (New England Biolabs). After sorting, plates were spun down and frozen at −80 °C.

Libraries were prepared as described previously (Ziegenhain et al. 2017; Soumillon et al. 2014). Briefly, proteins were digested with Proteinase K (Ambion) followed by desiccation to inactivate Proteinase K and reduce the reaction volume. RNA was then reverse transcribed in a 2 μL reaction at 42°C for 90 min. Unincorporated barcode primers were digested using Exonuclease I (Thermo Fisher). cDNA was pooled using the Clean & Concentrator-5 kit (Zymo Research) and PCR amplified with the KAPA HiFi HotStart polymerase (KAPA Biosystems) in 50 μL reaction volumes.

### mcSCRB-seq cDNA synthesis

Cells were dissociated using trypsin and resuspended in PBS. Single cells (“3 drops” purity mode) were sorted into 96-well DNA LoBind plates (Eppendorf) containing 5 μl lysis buffer using a Sony SH800 sorter (Sony Biotechnology; 100 μm chip). Lysis buffer consisted of a 1:500 dilution of Phusion HF buffer (New England Biolabs), 1.25 μg/μl Proteinase K (Clontech) and 0.4 μM barcoded oligo-dT primer (E3V6NEXT, IDT). After sorting, plates were immediately spun down and frozen at −80 °C. For libraries containing ERCCs, 0.1 μl of 1:80,000 dilution of ERCC spike-in Mix 1 was used.

Before library preparation, proteins were digested by incubation at 50 °C for 10 minutes. Proteinase K was then heat-inactivated for 10 minutes at 80 °C. Next, 5 μl reverse transcription master mix consisting of 20 units Maxima H- enzyme (Thermo Fisher), 2x Maxima H- Buffer (Thermo Fisher), 2 mM each dNTPs (Thermo Fisher), 4 μM template-switching oligo (IDT) and 15% PEG 8000 (Sigma-Aldrich) was dispensed per well. cDNA synthesis and template-switching was performed for 90 minutes at 42 °C. Barcoded cDNA was then pooled in 2 ml DNA LoBind tubes (Eppendorf) and cleaned-up using SPRI beads. Purified cDNA was eluted in 17 μl and residual primers digested with Exonuclease I (Thermo Fisher) for 20 min at 37 °C. After heat-inactivation for 10 min at 80 °C, 30 μl PCR master mix consisting of 1.25 U Terra direct polymerase (Clontech) 1.66x Terra direct buffer and 0.33 μM SINGV6 primer (IDT) was added. PCR was cycled as given: 3 min at 98 °C for initial denaturation followed by 15 cycles of 15 sec at 98°, 30 sec at 65 °C, 4 min at 68 °C. Final elongation was performed for 10 min at 72 °C.

### Library Preparation

Following preamplification, all samples were purified using SPRI beads at a ratio of 1:0.8 with a final elution in 10 μL of H_2_O (Invitrogen). The cDNA was then quantified using the Quant-iT PicoGreen dsDNA Assay Kit (Thermo Fisher). Size distributions were checked on High-Sensitivity DNA chips (Agilent Bioanalyzer). Samples passing the quantity and quality controls were used to construct Nextera XT libraries from 0.8 ng of preamplified cDNA.

During library PCR, 3’ ends were enriched with a custom P5 primer (P5NEXTPT5, IDT). Libraries were pooled and size-selected using 2% E-Gel Agarose EX Gels (Life Technologies), cut out in the range of 300-800 bp, and extracted using the MinElute Kit (Qiagen) according to manufacturer’s recommendations.

### Sequencing

Libraries were paired-end sequenced on high output flow cells of an Illumina HiSeq 1500 instrument. 16 bases were sequenced with the first read to obtain cellular and molecular barcodes and 50 bases were sequenced in the second read into the cDNA fragment. When several libraries were multiplexed on sequencing lanes, an additional 8 base i7 barcode read was done.

### Primary Data Processing

All raw fastq data was processed using zUMIs together with STAR to efficiently generate expression profiles for barcoded UMI data (Parekh et al. 2017; Dobin et al. 2013). For UHRR experiments, we mapped to the human reference genome (hg38) while mouse cells were mapped to the mouse genome (mm10) concatenated with the ERCC reference. Gene annotations were obtained from Ensembl (GRCh38.84 or GRCm38.75). Downsampling to fixed numbers of raw sequencing reads per cell were performed using the “-d” option in zUMIs.

### Filtering of scRNA-seq libraries

After initial data processing, we filtered cells by excluding doublets and identifying failed libraries. For doublet identification, we plotted distributions of total numbers of detected UMIs per cell, where doublets were readily identifiable as multiples of the major peak.

In order to discard broken cells and failed libraries, spearman rank correlations of expression values were constructed in an all-to-all matrix. We then plotted the distribution of “nearest-neighbor” correlations, ie. the highest observed correlation value per cell. Here, low-quality libraries had visibly lower correlations than average cells.

### Estimation of cellular mRNA content

For the estimation of cellular mRNA content in mouse ES cells, we utilized the known total amount of ERCC spike-in molecules added per cell. First, we calculated a “detection efficiency” as the fraction of detected ERCC molecules by dividing UMI counts to total spike ERCC molecule counts. Next, dividing the total number of detected cellular UMI counts by the “detection efficiency” yields the number of estimated total mRNA molecules per cell.

### ERCC Analysis

In order to estimate sensitivity from ERCC spike-in data, we modeled the probability of detection in relation to the number of spiked molecules. An ERCC transcript was considered as detected from 1 UMI. For each cell, we fitted a binomial logistic regression model to the detection of ERCC genes given their input molecule numbers. Using the MASS R-package, we determined the molecule number necessary for 50% detection probability.

For public data from *Svensson et al.*, we used their published molecular abundances calculated using the same logistic regression model obtained from “Supplementary Table 2” (https://www.nature.com/nmeth/journal/v14/n4/extref/nmeth.4220-S3.csv) (Svensson et al. 2017). For Quartz-seq2 (Sasagawa et al. 2017), we obtained expression values for ERCCs from Gene Expression Omnibus (GEO; GSE99866), sample GSM2656466; for Chromium (Zheng et al. 2017) we obtained expression tables from the 10x Genomics webpage (https://support.10xgenomics.com/single-cell-gene-expression/datasets/1.1.0/ercc) and for SCRB-seq, Smart-seq2, CEL-seq2/C1, MARS-seq and Smart-seq/C1 (Ziegenhain et al. 2017), we obtained count tables from GEO (GSE75790). For these methods, we calculated molecular detection limits given their published ERCC dilution factors.

### Power Simulations

For power simulation studies, we used the *powsimR* package (Vieth et al. 2017). Parameter estimation of the negative binomial distribution was done using scran normalized counts at 500,000 raw reads per cell (Lun, Bach, and Marioni 2016). Next, we simulated two-group comparisons with 10% differentially expressed genes. Log2 fold-changes were drawn from a normal distribution with mean of 0 and a standard deviation of 1.5. In each of the 25 simulation iterations, we draw equal sample sizes of 24, 48, 96, 192 and 384 cells per group and test for differential expression using ROTS (Seyednasrollah et al. 2015) and scran normalization (Lun, Bach, and Marioni 2016).

### Batch Effect Analysis

In order to detect genes differing between batches of one scRNA-seq protocol, data were normalized using scran (Lun, Bach, and Marioni 2016). Next, we tested for differentially expressed genes using limma-voom (Ritchie et al. 2015; Law et al. 2014). Genes were labelled as significantly differentially expressed between batches with Benjamini-Hochberg adjusted p-values < 0.01.

## Author contributions

CZ & WE conceived the study. JWB, CZ, AJ & LEW performed experiments and prepared sequencing libraries. JG & JWB cultured mouse ES cells. Sequencing data was processed by SP & CZ. JWB, CZ, AJ & BV analyzed the data. JWB, CZ, AJ, IH & WE wrote the manuscript.

## Acknowledgments

We thank Ines Bliesener for expert technical assistance. We are grateful to Magali Soumillon and Tarjei Mikkelsen for providing the original SCRB-seq protocol and to Stefan Krebs and Helmut Blum for sequencing.

This work was supported by the Deutsche Forschungsgemeinschaft (DFG) through LMUexcellent and the SFB1243 (Subproject A14/A15).

## Data availability

RNA-seq data generated here are available at GEO under accession GSE103568. Analysis code to reproduce all main Figures can be found at: https://github.com/cziegenhain/Bagnoli2017.

